# Outbreeding depression as a selective force on mixed mating in the mangrove rivulus fish, *Kryptolebias marmoratus*

**DOI:** 10.1101/2021.02.22.432322

**Authors:** Jennifer D. Gresham, Kever A. Lewis, Stephanie P. Summers, Percy E. Gresham, Ryan L. Earley

**Affiliations:** Emory University; University of Alabama; University of Alabama at Birmingham

**Author notes:** Corresponding author: Jennifer D. Gresham.

## Abstract

Mixed mating, a reproduction strategy utilized by many plants and invertebrates, optimizes the cost to benefit ratio of a labile mating system. One type of mixed mating includes outcrossing with conspecifics and self-fertilizing one’s own eggs. The mangrove rivulus fish (*Kryptolebias marmoratus)* is one of two vertebrates known to employ both self-fertilization (selfing) and outcrossing. Variation in rates of outcrossing and selfing within and among populations produces individuals with diverse levels of heterozygosity. I designed an experiment to explore the consequences of variable heterozygosity across four ecologically relevant conditions of salinity and water availability (10‰, 25‰, and 40‰ salinity, and twice daily tide changes). I report a significant increase in mortality in the high salinity (40‰) treatment. I also report significant effects on fecundity measures with increasing heterozygosity. The odds of laying eggs decreased with increasing heterozygosity across all treatments, and the number of eggs laid decreased with increasing heterozygosity in the 10‰ and 25‰ treatments. Increasing heterozygosity also was associated with a reduction liver mass and body condition in all treatments. My results highlight the fitness challenges that accompany living in mangrove forests ecosystem and provide the first evidence for outbreeding depression on reproductive and condition-related traits.

## Introduction

Most eukaryotic organisms engage in sexual reproduction and, in most vertebrates, there are no alternatives to sex. Despite the prevalence of sex, sixty years of modeling suggests that it should be difficult to sustain over evolutionary time, and has remained a fundamental question for biologists for well over 100 years (Darwin 1876; Maynard Smith 1971; Bell 1982; Hartfield and Keightley 2012). A mixed mating strategy, whereby individuals can either fertilize their own eggs (i.e., self-fertilization; selfing) or mate with a conspecific, eliminates some of the costs of traditional sexual reproduction. With self-fertilization, individuals avoid any energy use and risk associated with searching for a mate (Lehtonen et al. 2012); can overcome situations where there might be a lack of mates or pollinators (Baker 1955, reviewed in Pannell et al. 2015); and do not have to contend with diluting well-adapted genotypes (Allard et al. 1972; Becks and Agrawal 2012). However, the opportunity to engage in traditional mating allows species with mixed mating systems to also exploit the benefits of sex, including an escape from inbreeding depression (Jennings 1914; Charlesworth and Charlesworth 1987) and the production of novel genotypes that might be better adapted to a fluctuating environment or at “outrunning” coevolving parasites (Muller 1932; Bell 1988; Lively 2010; Morran et al. 2011; Hartfield and Keightley 2012).

In species with a mixed mating strategy, rates of selfing and outcrossing are expected to vary as biotic (e.g., population density, parasite density, resource availability) or abiotic (e.g., temperature or precipitation variability) environmental conditions alter the cost-benefit ratio associated with each mode of reproduction (Jarne and Charlesworth 1993; Escobar et al. 2011). The economics of employing selfing versus outcrossing as the predominant mode of reproduction can vary spatially. For instance, at the range edge where population density is low and mates are limited, selection should favor individuals that reproduce by selfing (Baker 1955; Pannell 2000; Grossenbacher et al. 2015). In environments that are stochastic or otherwise challenging, selection should favor recombination of alleles and genetically diverse offspring, and thus individuals that reproduce by outcrossing (Fisher 1930; Muller 1932; Williams 1975). The assumption is that some genetically diverse progeny are more likely to survive in challenging conditions, although models and empirical data do not always support this assumption (Hines and Moore 1981; Robson et al. 1999; Cheptou and Schoen 2002). Rates of selfing versus outcrossing also can vary temporally. Mixed mating should be evolutionarily stable when selection regimes fluctuate between those that would favor selfing and those that would favor outcrossing thus, maintaining both selfing and outcrossing alleles in the population (Goodwillie et al. 2005).

Importantly, for a mixed mating strategy to be evolutionarily stable, natural selection must favor individuals that are able to accurately assess environmental cues and adopt the strategy (selfing or outcrossing) that affords highest fitness for themselves and their offspring (Bull 1981; Bell 1982; Shuster 2008). Many hermaphroditic plants preferentially outcross when and where pollinators and conspecifics are available, but self when these mating opportunities are scarce (Baker 1955; Kalisz et al. 2004; Busch 2005; Hesse and Pannell 2011; de Waal et al. 2014). Similarly, some snails and cestodes will delay selfing until late in life after waiting for potential mates (Jordaens et al. 2007; Escobar et al. 2011). Parasite prevalence also influences mating strategy. Studies using *Caenorhabditis elegans* as the host organism have revealed that novel bacteria strains induce outcrossing compared to strains with which the nematode worms have coevolved, and that the rate of outcrossing compared to selfing also depended on the virulence of the bacteria strain (Lynch et al. 2018).

Often, the costs of selfing and outcrossing do not become evident until individuals are exposed to environmental challenge(s). For instance, obligate outcrossing species may only experience costs associated with finding mates when at the range edge or when mating opportunities are limited for other reasons (Igic and Busch 2013). Obligate outcrossing can also break up well-adapted genotypes, presenting a cost in stable environments. Rotifers increase the frequency of sexual reproduction (outcrossing) while adapting to a new food environment but decrease the frequency of outcrossing once adaptation to the stable environment has been established and sexually-derived offspring become less fit than their asexual congeners (Becks and Agrawal 2012). These results are expected to be similar when comparing obligate outcrossing and obligate selfing populations of the same species. Indeed, higher and less variable fitness was found in the progeny of selfing lineages of the fungus *Aspergillus nidulans* compared to outcrossing lineages when the environment was held constant, even though the parent populations had equal fitness, an example of outbreeding depression (López-Villavicencio et al. 2013). The costs of selfing, not being able to quickly adapt to a new abiotic environment or new parasites, can lead to extinction (i.e., the “dead end hypothesis”), because successive generations are increasingly parasitized and have lower fitness than previous generations (Stebbins 1957; Lande and Schemske 1985; Busch and Delph 2017). Experimental coevolution of the *C. elegans* host with *Serratia marcesans* bacteria conclusively demonstrated that obligate selfing populations were quickly driven to extinction, while outcrossing populations endured (Morran et al. 2011). Models indicate that the lack of adaptive potential, which can lead to extinction in selfing populations, requires a very specific set of genetic and environmental conditions. Burgarella et al. (2015) showed empirically that selfing populations of freshwater snails (*Galba truncatula*) have avoided the consequences of inbreeding depression and probable extinction by escaping interspecific competition in their unstable environments. For these reasons, it is important to assess inbreeding and outbreeding depression using multiple measures, across multiple environments.

Species that can alternate between selfing and outcrossing offer unique insights into the costs and benefits of both strategies. These species allow comparisons of strategies across different lineages while eliminating the complications of phylogeny, such as different evolutionary histories and adaptations to different environments. With some species, we can often control for, or exploit, population differences in the extent of selfing and outcrossing, while also measuring inbreeding and/or outbreeding depression across multiple environments. However, empirical studies that document the costs and benefits of mixed mating strategies within a species are limited to a few model organisms (*Leavenworthia alabamica;* Busch et al. 2010, Layman et al. 2017*, C. elegans*; Anderson et al. 2010, Penley et al. 2018, and some pulmonates; reviewed in Jordaens et al. 2007). Here, I present results of an experiment using an emerging model for the evolution of mating strategies, one of two vertebrates that can toggle between self-fertilization and outcrossing as modes of reproduction. This experiment was designed to measure indicators of inbreeding and/or outbreeding depression (proxy measures for understanding the relative costs of selfing and outcrossing, respectively) across four different environmental conditions in the mangrove rivulus fish (*Kryptolebias marmoratus,* hereafter “rivulus”). Populations of rivulus consist of simultaneous hermaphrodites, a small but variable proportion of males, and no females (androdioecy). Hermaphrodites reproduce primarily by self-fertilization, but occasionally outcross with males. Rates of outcrossing within a population correlates with the proportion of males, and both vary greatly across rivulus’ range (Mackiewicz et al. 2006; Tatarenkov et al. 2009). It is likely, but not experimentally documented, that there is also temporal variation in rates of outcrossing within populations. This mixed mating strategy is stable and has persisted for hundreds of thousands of years (Sato et al. 2002; Tatarenkov et al. 2009), leading to the conclusion that both outcrossing, which increase offspring heterozygosity, and selfing, which decreases offspring heterozygosity, offer important benefits. To better understand how mixed mating has endured in this species, I designed an experiment to measure the effects of heterozygosity on fitness across four different ecologically relevant environmental treatments. I hypothesized that one or more fitness characteristics would be affected by heterozygosity. Specifically, I predicted that some heterozygosity would afford better fitness than complete homozygosity (inbreeding depression). I also predicted that inbreeding depression would become most evident in stressful environments.

## Methods

### Experimental Design

This experiment was designed to measure fitness-related traits that might indicate either inbreeding or outbreeding depression in rivulus. 1098 fish representing 69 genetically distinct lineages, with varying levels of heterozygosity, were divided among four ecologically relevant treatments: control (constant volume of 25‰ water), low salinity (constant volume of 10‰ water), high salinity (constant volume of 40‰ water), and tidal (varying volume of 25‰ water). A summary of information for each of the 69 lineages and their distribution across treatments can be found in Appendix S1: Table S1.1.

The sample of experimental individuals were two or three generations removed from their wild caught progenitors to minimize parental effects. The progenitors of these experimental fish were held individually in 750 ml clear plastic containers (Rubbermaid^®^ Take-A-Long Deep Squares) filled with approximately 600 mL of 25‰ synthetic salt water prepared with Instant Ocean^®^ synthetic salt. Each container had a large marble-sized piece of Poly-Fil 100% polyester fiber stuffing (Poly-Fil^®^ Premium Polyester Fiber Fill) suspended from the lid, which was used as a reproductive substrate to induce egg laying and enable egg collection. Each progenitor fish was fed ~2000 *Artemia sp.* nauplii suspended in 4 mL of 25‰ water daily between 0900-1700 h. Eggs from the progenitor fish were collected once per week by manually separating the eggs from the Poly-Fil substrate. The eggs were put into plastic 60 mL cups (shipyouraquatics.com, Item SYAPC400500) filled with approximately 6 mL of 25‰ synthetic salt water, prepared with Instant Ocean^®^ synthetic salt, until hatching. The progenitors and experimental fish were held in the same room with 12h light:12h dark photoperiod and temperature maintained at 25.5 ± 0.59°C (mean ± SEM).

At hatching, experimental fish were randomly assigned to one of the four treatments using a random number generator. At least four fish per lineage were placed in each treatment. Lineages were chosen from two regions of rivulus’ range, Belizean Cays and Florida Keys, with the goal of maximizing variation in heterozygosity available in our colony. Previously, the heterozygosity of wild caught fish (F0 generation) was measured using 32 microsatellites (Mackiewicz et al. 2006) and our colony contains fish ranging from 0 to 23 heterozygous microsatellite loci. On average, heterozygosity should be reduced by one half each generation. This likely reduced the heterozygosity range to between 0 and 6 heterozygous microsatellites in my experimental generation but was not confirmed by microsatellite analysis.

The experimental phase for each individual fish was divided into two periods. The first period (juvenile) was from 0 to 89 days post hatch (dph). The second period (treatment) was from 90 to 180 dph. During the juvenile period, individual fish were held in 473 mL white plastic cups with lids (VWR Multipurpose Containers; 473 mL Translucent HDPE with snap lid), filled with approximately 400 mL of 25‰ salt water prepared with Instant Ocean® synthetic salt. This water was replaced every 30 days, during which time the fish was measured for mass and length (described below). At 89 dph, the fish was transferred into the same type of 473 mL white plastic cup, but with 8, 0.36 cm (9/64” drill bit) diameter holes drilled around the bottom edge to allow for water flow. The fish and cup were then placed into the appropriate transition tank for the individual’s assigned treatment. At 90 dph, each fish was measured for standard length and mass, then transferred to its treatment tank (still in its cup with drilled holes; treatments were control, low salinity, high salinity, tide as described below) for 90 more days or until death, whichever happened first. Every 30 days, each fish was removed from its cup, measured for standard length and mass, and assessed for morphological changes indicating sex change. To measure fish for standard length, the fish was placed on a small plastic metric ruler and photographed with a smartphone. Photos were then uploaded into ImageJ and fish were measured digitally; standard length was taken as the distance between the tip of the nose and the posterior margin of the caudal peduncle. Mass was obtained by dabbing the fish on a clean paper towel to remove excess water, placing it into two small plastic weigh boats taped together to form a “clam” (to prevent the fish from jumping out), and weighed on a digital balance tared to the mass of the weigh boats. Fish were assessed for the morphological characters that accompany sex change (orange freckles or orange skin) as described in Scarsella et al. (2018). At 90 dph, a large marble-sized ball of synthetic poly-fiber was suspended from the top of each cup to create an egg-laying substrate. Throughout the treatment period, each cup was checked for eggs once every seven days. These eggs were collected into 60 mL portion cups containing water with the same salinity from which the eggs were collected. Eggs were viewed under a stereomicroscope to score each as fertilized, unfertilized, or dead. Fertilized eggs are identifiable by the perivitelline membrane inside the chorion; unfertilized eggs do not have the perivitelline membrane (Harrington 1963, Sucar et al. 2016). Dead eggs are identifiable because they are opaque and range in color from white to red. Fertilized eggs were allowed to hatch and hatchlings were euthanized by immersion in 4°C water and measured as described above. The treatment period ended at 180 dph. On this day, surviving fish were removed from treatment tanks, euthanized by immersion in 4°C water, measured as previously described, and dissected. The liver was removed and weighed. Gonads were confirmed as either an ovotestis (hermaphrodite) or testis (male), removed, and weighed. During the experiment, fish were fed ~1000 *Artemia sp.* nauplii suspended in 2 mL of water at the appropriate salinity once per day; fish 0 – 30 dph were fed ~500 nauplii suspended in 1 mL of water at the appropriate salinity.

### Treatment Construction

Transition and treatment tanks were constructed from 53 L Sterilite storage containers with a raised false floor made from black corrugated plastic sitting atop a cement block. The white fish cups sat on top of this corrugated plastic. Each transition tank consisted of a single container. The transition tank for the low salinity treatment contained a constant volume of water at 17‰. The transition tank for high salinity contained a constant volume of water at 32‰. Because fish moving into the control or tide treatments were not changing salinity, they shared a “transition” tank of a constant volume of water at 25‰.

Each treatment tank consisted of two storage containers connected via 1.9 cm PVC pipe and water tight bulkheads (Bulkreefsupply.com, 1.9 cm ABS thread x thread); this was considered to be one dual tank. There were two separate dual tanks for each treatment (this is the random “tank” variable in the statistical models described below). Both transition and treatment tanks received a constant (except the tide treatment) inflow of water from individual sumps; water was pumped in over the side wall using Hydor Centrifugal pumps and flexible plastic tubing. Water from the tanks drained back into the sumps via flexible plastic tubing connected to a bulkhead placed 2.54 cm from the top of the tank. The height of the drain tube and water pressure from the sump was specifically determined to keep the white fish cups filled to 600 mL. Each sump also acted as a filter and contained white filter felt (pentairaes.com, Part no. PF11P), a large sachet filled with carbon pellets (Marineland^®^ Black Diamond Activated Carbon Filter Media), and about 13 cm of plastic biomedia (Sweetwater^®^ SWX Bio-Media) from top to bottom. The control, high salinity, and low salinity treatments consisted of continuously flowing water at 25‰, 40‰, and 10‰, respectively. Although water was continuously flowing through the sumps and tanks, the current within the cups was just enough to remove wastes and maintain a constant salinity and temperature, as checked with dye tests prior to the beginning of the experiment. The tide treatments had continuously flowing water at 25‰ for two-5.5 hour periods (high tide) per day. High tide periods were interspersed with low tides, which consisted of 5.5 hour intervals where the tank water was drained so that each fish cup contained 10 mL of still water. Water levels were controlled by the sump pump connected to digital timers and an extra tank pump connected to digital timers (Defiant^®^, 15 Amp 7-Day Indoor Plug-In Digital Timer with 2-Grounded Outlets). It took 30 minutes to transition from high tide to low tide and vice versa. Water temperature was maintained at 23.2 ± 0.03 °C (mean ± SEM). Lighting was automatically controlled to create a 12 h light:12 h dark photoperiod. Figures of the treatment tanks are available in Gresham et al. (2020, Chapter 2). The entire experimental period lasted 17.5 months. The University of Alabama Institutional Animal Care and Use Committee approved all procedures (IACUC Protocol # 15-04-0102).

### Fitness Endpoints and Statistical Analyses

Survival to 180 dph as a binary response (yes or no) and age at death were recorded. To understand whether heterozygosity affected the probability of surviving the juvenile period, I used a Cox proportional hazards model with age of death as the response and heterozygosity as the predictor, censored by whether the fish survived the juvenile period (yes = 1, no = 0). To understand how heterozygosity and treatment affected the probability of surviving the treatment period, I used a Cox proportional hazards model with age of death as the response and with heterozygosity, treatment, and their interaction as predictors, censored by whether the fish survived the treatment period (yes = 1, no = 0).

Mass and standard length were measured every 30 days, with Fulton’s K body condition ([mass/standard length^3^] x 100) calculated from these measurements. I used general linear mixed models to examine how size traits were affected by heterozygosity and/or treatment. Juvenile period models included heterozygosity as the only predictor of body size. Treatment period models included heterozygosity, treatment, and the heterozygosity x treatment interaction as fixed predictors, and tank as a random effect.

I quantified various aspects of fecundity. Egg laying as a binary response (yes = 1, no = 0) was recorded, as well as the total number of eggs, and the total number of fertilized, unfertilized, and dead eggs laid by each fish. Hatch success of fertilized eggs collected in the treatments was also recorded. To understand how heterozygosity and treatment affected fecundity, I ran multiple models with heterozygosity, treatment, and their interaction as fixed predictors and tank as a random effect. To determine the significance of treatment and heterozygosity (and the interaction) on whether or not individuals laid eggs, I ran a general linear model with a binomial distribution and logit link function, and calculated odds ratios where appropriate. I also ran generalized regression models, with a zero-inflated Poisson distribution and maximum likelihood parameter estimation, for number of eggs laid and the proportion of eggs that were unfertilized. I can infer how the proportion of fertilized eggs is affected by heterozygosity and treatment from the unfertilized egg model. Finally, I calculated hatch success as the proportion of fertilized eggs that successfully hatched for each individual. A general linear mixed model was run as described above, with hatch success as the response variable.

At the end of the treatment period, surviving fish were dissected, and the livers and gonads resected and measured for mass. To determine how heterozygosity and treatment affected gonad size, I calculated residual gonad mass from a regression of body mass versus gonad mass. I ran a general linear mixed model with gonad residuals as the response variable; heterozygosity, treatment, heterozygosity x treatment as the fixed effects, and tank as a random effect. I analyzed the effects of heterozygosity, and treatment on liver size in the same manner as gonad mass, with liver residuals calculated and used as the response variable. I also recorded which individuals changed sex to male and reported those results in Gresham et al. (2020, Chapter 2).

When appropriate, multiple comparisons were conducted using Tukey’s HSD pairwise comparisons, which controls for compounding Type I error. Although 872 individuals survived the juvenile period and 675 individuals survived the treatment period, degrees of freedom in the models will vary slightly as reliable measurements were not obtained for all endpoints in all individuals. Size and survival analyses were run using JMP Pro version 15.0 (JMP^®^, Version 15 Pro 2019). All fecundity models were analyzed in RStudio, using the *car*, *lme4*, *emmeans*, and *glmmTMB* packages (Fox and Weisberg 2011; Bates et al. 2015; RStudioTeam 2016; Brooks et al. 2017; R Core Team 2018; Lenth 2019).

## Results

Survival to 90 dph was not affected by individual heterozygosity and survival from 90 to 180 dph was affected only by treatment (Table 1.1). Individuals in high salinity had a significantly lower probability of surviving the treatment period than those in low salinity, tide or control conditions; none of the other comparisons were statistically significant (Table 1.1, Figure 1.1).

**Table 1.1.**
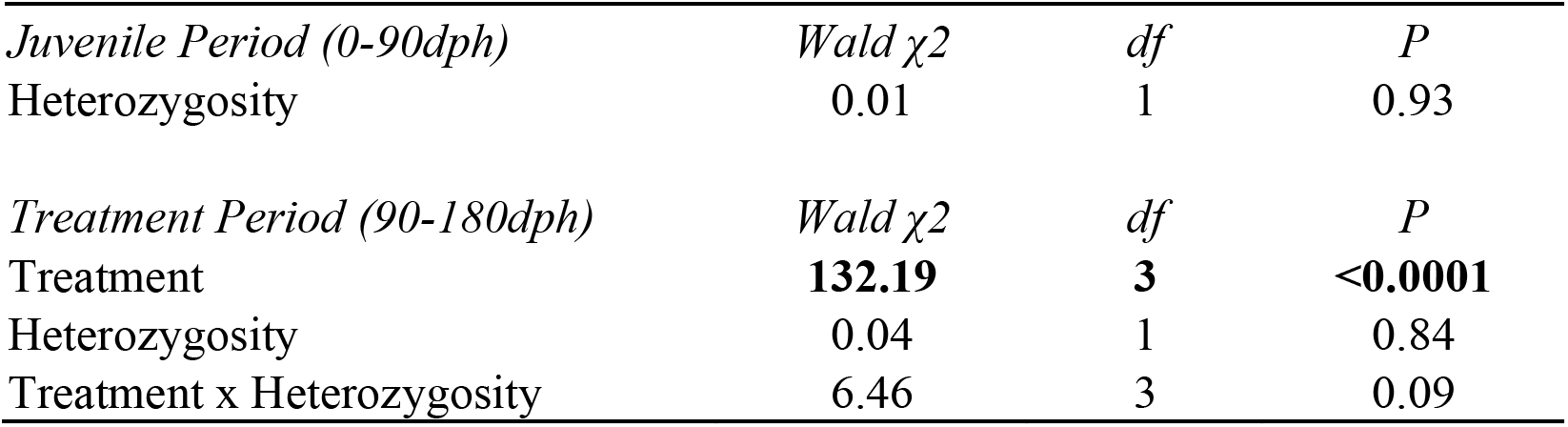
Summary of Cox proportional hazard models of survival to the end of the juvenile period and the end of the treatment period. All juveniles were kept in the same constant condition of 25‰. There were four treatments in the treatment period (control, low salinity, high salinity, and tide). Heterozygosity was modeled as a continuous variable ranging from 1 - 6 heterozygous microsatellites (out of 32). Significant effects are shown in bold. Whole model test data and the parameter estimates can be found in the appendix, Table S1.1. P = P-value, df = degrees of freedom.

| *Juvenile Period (0-90dph)* | *Wald $\chi^2$* | *df* | *P* |
| --- | --- | --- | --- |
| Heterozygosity | 0.01 | 1 | 0.93 |
| *Treatment Period (90-180dph)* | *Wald $\chi^2$* | *df* | *P* |
| Treatment | **132.19** | **3** | **<0.0001** |
| Heterozygosity | 0.04 | 1 | 0.84 |
| Treatment x Heterozygosity | 6.46 | 3 | 0.09 |

**Figure 1.1:**
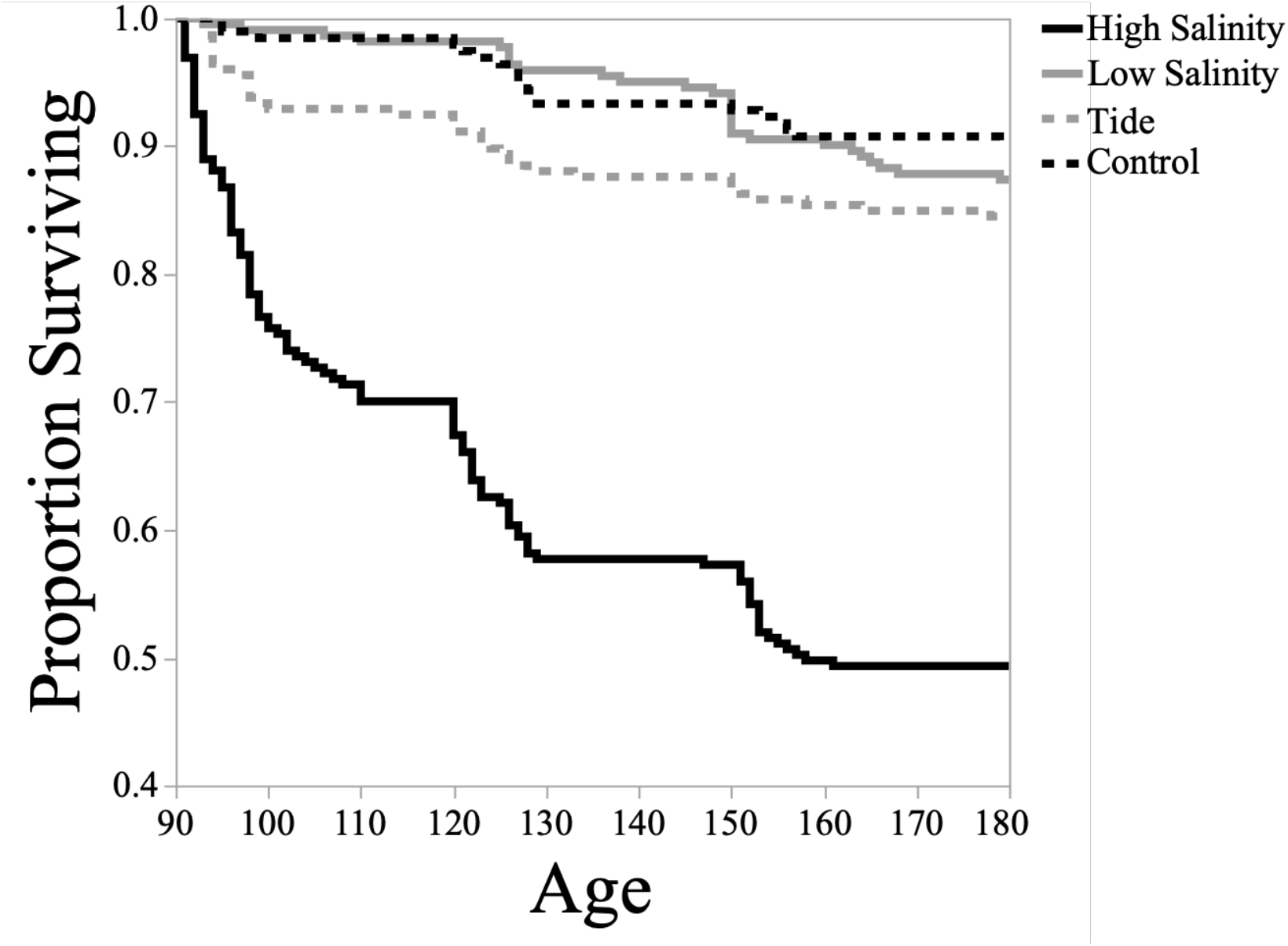
Treatment effects on survival after fish entered treatment at 90 dph. Survival in high salinity was significantly lower than all other treatments. There were not differences among the remaining three treatments.

At the end of the juvenile period, there was a marginal effect of heterozygosity on mass and a significant effect of heterozygosity on standard length; animals with greater heterozygosity tended to be heavier and were significantly longer than their more homozygous counterparts (Appendix S1: Table S1.2). Juvenile body condition (Fulton’s K), however, did not vary with heterozygosity (Appendix S1: Table S1.2). The treatment in which the animals lived from 90-180 dph significantly affected mass (Figure 1.2), standard length, and body condition (Appendix S1: Figure S1.1). Individuals in the tide treatment were significantly lighter, shorter, and were in worse body condition than individuals in all other treatments. Individuals in low salinity also were significantly heavier, but not significantly longer or in better condition, than those in the control treatment (Figure 1.2; Appendix S1: Figure S1.1). Body condition decreased with increasing heterozygosity, irrespective of treatment (Appendix S1: Figure S1.1).

**Figure 1.2:**
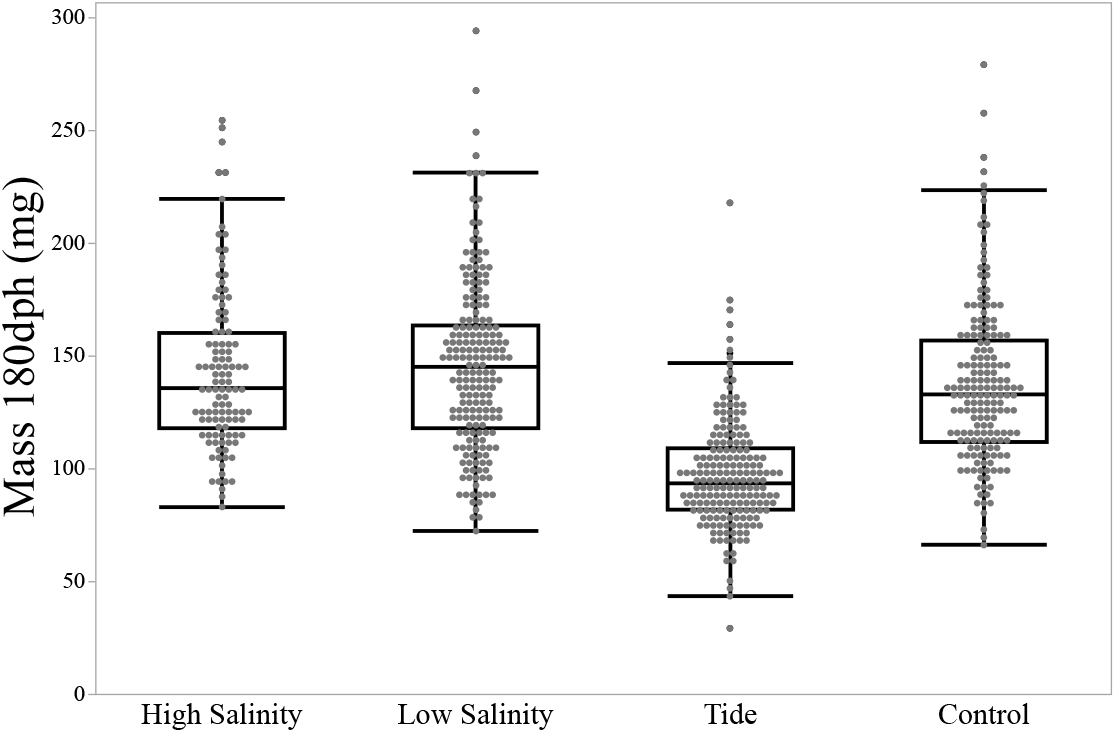
Treatment effects on mass at 180 dph. Fish in the tide treatment were significantly lighter and shorter and had lower body condition values than fish in all other treatments. Body condition also decreased with increasing heterozygosity in all treatments. Graphs for length and body condition are available as Supplement Figure S1.1. Tables describing treatment effects on all size characters are available in the Supplement as Figure S1.1 and Table S1.2.

Whether or not an individual laid any eggs was affected, independently, by treatment and heterozygosity (Table 1.2). Animals in the control treatment were 4-5 times more likely to lay eggs, while animals in low salinity were 2.5-3 times more likely to lay eggs, than those in the high salinity and tide treatments (Figure 31.A; Appendix S1: Table S1.3). The likelihood of laying eggs decreased with increasing heterozygosity across treatments (Table 1.2). Both treatment and heterozygosity significantly affected the number of eggs laid, and the relationship between heterozygosity and fecundity varied with treatment (i.e., significant treatment x heterozygosity interaction) (Table 1.2). In both control and low salinity, the number of eggs laid decreased with increasing heterozygosity, a trend that was absent in both high salinity and tide treatments (Figure 1.3B). Heterozygosity also affected the proportion of fertilized/unfertilized eggs in a treatment-dependent fashion. In both high and low salinity, the proportion of fertilized eggs increased with increasing heterozygosity; but in the control, the proportion of fertilized eggs decreased with increasing heterozygosity. In the tide treatment, virtually all eggs were fertilized (98.9 ± 4.7%) (Appendix S1: Table S1.4 and Figure S1.2). Hatch success was not affected by treatment, heterozygosity, or their interaction (Appendix S1: Table S1.4).

**Figure 1.3.**
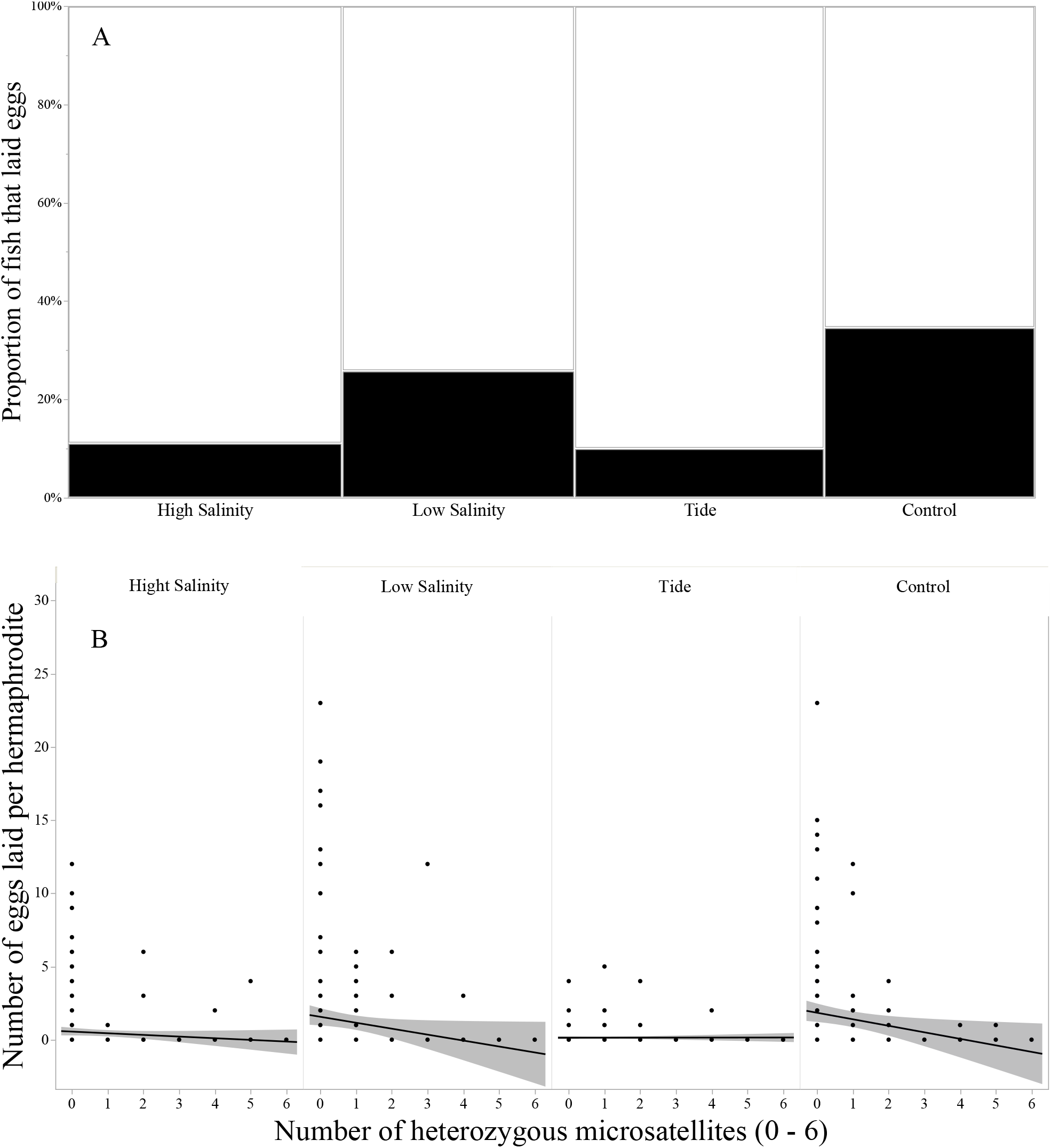
Treatment and heterozygosity effects on fecundity traits. A) The proportion of fish that laid any eggs was only significantly affected by treatment. Fish in low salinity and control treatments were more likely to lay eggs than those in tide and high salinity. B) The number of eggs laid was affected by an interaction of treatment and heterozygosity. In both low salinity and control, the number of eggs laid decreased as function of increasing heterozygosity. This trend is absent in both high salinity and tide. Whole model test data and the parameter estimates can be found in the appendix, Table S1.3.

**Table 1.2.** Summary of generalized mixed models for whether or not hermaphrodites laid eggs and the number of eggs laid by hermaphrodites. There were four treatments (control, low salinity, high salinity, and tide). Heterozygosity was modeled as a continuous variable ranging from 1 - 6 heterozygous microsatellites (out of 32). Significant effects are shown in bold. Whole model test data and the parameter estimates can be found in the appendix, Table S1.3. P = P-value, df = degrees of freedom, Y = yes, N = no.

| *Nominal Logistic for Laying Eggs, Y/N* | *Wald $\chi^2$* | *df* | *P* |
| --- | --- | --- | --- |
| Treatment | **35.73** | **3** | **<0.0001** |
| Heterozygosity | **3.87** | **1** | **0.049** |
| Treatment x Heterozygosity | 2.84 | 3 | 0.42 |
| *Number of eggs laid* | *Wald $\chi^2$* | *df* | *P* |
| Treatment | **28.43** | **3** | **<0.0001** |
| Heterozygosity | **35.26** | **1** | **<0.0001** |
| Treatment x Heterozygosity | **19.93** | **3** | **0.0002** |

Residual gonad mass differed significantly among treatments (Appendix S1: Table S1.5), a result driven largely by the fact that residual gonad mass was significantly lower in control than in low salinity animals; all other comparisons were non-significant. Heterozygosity was the only significant predictor of residual liver mass, which decreased as heterozygosity increased (Appendix S1: Table S1.5).

## Discussion

Understanding the evolution and maintenance of mating strategies requires that we know the costs and benefits of alternative strategies across the range of ecologically relevant environments. Mangrove rivulus fish employ a mixed mating strategy of self-fertilization and outcrossing, rates of which vary greatly across the species’ geographical range (Turner et al. 1992; Tatarenkov et al. 2009). To ascertain how heterozygosity affects fitness, and which environmental factors might determine the consequences of outcrossing and selfing, I designed an experiment to measure fitness differences among individuals with varying levels of heterozygosity in four different environments. I hypothesized that there would be fitness differences with varying levels of heterozygosity and predicted that individuals with less heterozygosity would show evidence of inbreeding depression. I also predicted that inbreeding depression might not be revealed until the individuals encountered challenging environments. Previously, Ellison et al. (2011) reported that parasite load increased as a function of decreasing heterozygosity in rivulus, indicating potential inbreeding depression in animals caught in the field. In other animals, studies have indicated that inbred individuals do not exhibit fitness decrements until presented with suboptimal ecological conditions. For instance, inbreeding experiments with *Drosophila melanogaster* showed a decrease in reproductive fitness in inbred lineages when exposed to temperature or metal stress, but not under control conditions (Miller 1994). Studies in species of conservation concern have shown little evidence of inbreeding depression under controlled conditions, but significant decreases in reproductive fitness once inbred animals are exposed to food shortages or competition (Galapagos finches; Keller et al. 2002, Mexican wolves, Fredrickson et al. 2007). In light of these studies, I constructed four ecologically relevant treatments: 25‰ salinity as my control; 10‰ and 40‰ salinity treatments as osmoregulatory stressors, and a twice daily low tide of 25‰ water to serve as a water availability stressor. To understand fitness consequences as best I could, I measured standard length, mass, body condition, fecundity, hatch success, incidence of sex change, liver and gonad masses, and mortality of all individuals. When there was a significant effect of heterozygosity, it always pointed to outbreeding depression. Ecological treatment also had profound effects on fitness characters, emphasizing the challenges associated with rivulus’ habitat. Treatment and heterozygosity also frequently interacted to explain variation in fitness.

Heterozygosity was associated with size in opposite ways during the juvenile and treatment periods, similar to what was documented previously in blue tits (*Cyanistes caeruleus*), which exhibited outbreeding depression in early life traits (e.g., decreased hatching rate), but inbreeding depression on late life traits (e.g., adult survival and reproduction; Olano-Marin et al. 2011). Here, I found that standard length and mass increased as a function of increasing heterozygosity during the juvenile period, without any variation in body condition. However, in the treatment period (> 90 dph), mass and length were not affected by heterozygosity, but body condition decreased with increasing heterozygosity.

I found evidence of outbreeding depression during the treatment period for traits associated with reproductive fitness. In all treatments, the odds that hermaphrodites would lay eggs decreased with increasing heterozygosity, and more heterozygous individuals laid proportionally fewer fertilized eggs. Further, in both the control and low salinity treatments, the absolute number of eggs laid by individuals decreased with increasing heterozygosity. This trend was absent in the high salinity and tide treatments, possibly because so few eggs were laid in those treatments. Individuals in the low salinity or control treatments were indeed significantly more likely to lay eggs than those in either the tide or high salinity treatments. Outbreeding depression related to reproduction is not uncommon in species with mixed mating systems. For instance, in *Leptosiphon jepsonii*, an annual herb that utilizes mixed mating, outcrossing populations had a lower seed set than selfing populations (Goodwillie and Knight 2006). The same has been found in Alabama gladecress (*Leavenworthia alabamica,* Busch et al. 2010). However, a meta-analysis by Winn et al. (2011), indicated that plants that employ mixed mating have as much inbreeding depression related to reproduction as purely outcrossing species. These contradictory conclusions make it much harder to find clear patterns with respect to the consequences of mixed mating and demonstrate the importance of studying as many species as possible.

In support of my hypothesis, I did find significant treatment differences for other fitness characters I measured, other than hatch success. There was significantly lower survival in high salinity compared to all other treatments. Lin and Dunson (1999) also showed more mortality in their 40‰ salinity treatment than in 12‰ or 1‰, suggesting that while 40‰ is an ecologically relevant salinity, it can have severe consequences on individual fitness and population size. Fitness consequences in higher salinities are further supported by rivulus’ preference to inhabit and lay eggs in lower salinity (5‰), as well as the increased hatching rate at low salinities (McCain et al. 2020). In addition to reproduction, size also was treatment-dependent. Individuals in the tidal treatment were significantly shorter and lighter than animals in all other treatments. Individuals in the low salinity treatment also were heavier than those in the control treatment, but not those in high salinity. Given considerable mortality in the high salinity treatment compared to the other treatments, and the absence of a size difference, I predicted *post hoc* that there might be selection against smaller animals in the high salinity treatment. I constructed proportional hazards models with age at death as the response and treatment, size (numerous variables run in separate models), and the interaction of treatment and size as fixed effects, censored by whether or not the individual lived to be 180 dph (yes = 1, no = 0; i.e., completed the treatment period). I modeled mass, standard length, and body condition at 90 dph (immediately before entering treatments) and 150 dph (the measurement prior to the end of study). Treatment and both mass and standard length significantly influenced whether individuals lived to 180 dph, but not body condition or the treatment by size interactions (Appendix S1: Table S1.6). I found that there was selection on smaller animals in all treatments (note that the negative parameter estimate in the model indicates that the hazard of mortality decreases with the increase of the independent variable). Animals that were lighter and shorter at 90 dph and 150 dph were less likely to survive the entire treatment period. Even for marine and euryhaline fish, high salinity can drive mortality and can even dictate sex ratios. The mangrove red snapper (*Lutjanus argentimaculatus*) suffers increased mortality during early life stages when exposed to higher salinity (40‰) than ocean salinity (32‰) ( Estudillo et al. 2000). On the contrary, the survival of sailfin mollies (*Poecilia latipinna*) varied between freshwater and saltwater conditions depending on the year and season (Trexler et al. 1992). High salinity during the juvenile period skews sexual differentiation significantly towards males in sea bass (*Dicentrarchus labrax*) (Saillant et al. 2003). There also is additional evidence that selection against small size is common in a variety of conditions. In the European sardine (*Sardina pilchardus*), there is evidence of higher mortality in smaller individuals in both the wild and laboratory conditions (Garrido et al. 2015). This has also been shown under multiple conditions in Baltic Cod (Grønkjær and Schytte 1999), two species of wrasses (Raventós and Macpherson 2005), brown demoiselle (Vigliola and Meekan 2002), and Atlantic cod (Meekan and Fortier 1996).

It should be noted that my animals were not given the option of escaping a bad environment (high salinity and tidal treatments), a contradiction to their behavior in the wild. Prior and current observations have shown that rivulus are comfortable out of water and have evolved multiple mechanisms to cope with low tide and high salinity conditions. Rivulus can restructure their gills and skin for aerial breathing (Ong et al. 2007; Turko et al. 2020), can coordinate multiple osmoregulatory responses to high salinity (Frick and Wright 2002; Sutton et al. 2018), and have means of escaping unfavorable conditions by slithering or jumping to alternate microhabitats (Taylor 2000; Pronko et al. 2013). Despite the fact that rivulus have mechanisms to adapt to non-optimal conditions such as the tide and high salinity treatments, it appears that living in such habitats over the long-term has significant fitness consequences.

I have shown that rivulus with higher heterozygosity experience outbreeding depression in body condition and most reproductive traits. However, I did not observe inbreeding or outbreeding depression in survival, hatch success, or other size traits (mass and standard length). However, this does not mean that heterozygosity is inconsequential for these other traits. The experimental individuals were two or three selfed generations removed from wild caught individuals. On average, for each generation of selfing, heterozygosity is reduced in each individual by half. Thus, heterozygosity differences among experimental individuals may not have been large enough to quantify fitness differences in survival, size, and/or hatchling success. The highest level of heterozygosity in my experimental animals may only be six out of thirty-two microsatellites. Previous reports about inbreeding and outbreeding depression have discussed the low likelihood of detecting inbreeding or outbreeding depression when the genome is already highly homozygous or heterozygous respectively, because additional homozygosity or heterozygosity is but a fraction of the entire genome (Szulkin and David 2011). I also could not measure the true heterozygosity of each experimental animal. While offspring of heterozygous parents experience an *average* one-half reduction in heterozygosity, it is not necessarily the same reduction of heterozygosity among all siblings. This source of error may have concealed inbreeding or outbreeding depression in this study because I was using expected instead of actual values for heterozygosity. Using microsatellites as a measure of heterozygosity also has limitations when assessing inbreeding and outbreeding depression. Hoffman et al. (2014) compared the incidence of lungworm infection in seals with heterozygosity measured with microsatellites and single nucleotide polymorphisms (SNPs), and found a highly significant correlation between lung infection and heterozygosity with SNP data, but only a marginally significant trend with microsatellite genotyping. In rivulus, Lins et al. (2018) reported a range of 0.0305% - 0.0554% SNP heterozygosity in individuals that were completely homozygous at 32 microsatellites, which is very small compared to outcrossing species but not zero.

The ability to observe inbreeding and outbreeding depression is very specific to the traits that are measured, and the conditions tested (Szulkin et al. 2010). I observed outbreeding depression in reproductive traits when the animals were exposed to abiotic stressors. There may also be inbreeding or outbreeding depression in other traits not measured (lifelong reproductive fitness, resource procurement), or in response to biotic stressors such as interspecific competition or parasites. Studies indicate a decrease in major histocompatibility complex (MHC) supertypes with increasing homozygosity, with the possible consequence of decreased immunity against a variety of parasites (Ellison et al. 2012). Indeed, Ellison et al. (2011) showed that more homozygous individuals were more susceptible to infections by multiple parasites. Molloy (2011) also reported that the size of rivulus males increases with increasing heterozygosity, potentially affecting their access to resources and mating opportunities. My results indicate that even moderate heterozygosity, that is much lower than we find in wild caught individuals (Tatarenkov et al. 2015), can cause significant fitness differences among individuals. This difference in reproductive fitness has the potential to drive evolution towards increased selfing rates but may be counterbalanced by other heterozygosity effects on fitness.

Understanding the how selection maintains a specific reproductive strategy requires assessing the costs and benefits of the strategy across multiple environments. In this study I measured nine fitness characters across four different environments in the mangrove rivulus fish, a vertebrate with a mixed mating strategy of selfing and outcrossing. In addition to demonstrating how challenging certain aspects of the mangrove habitats (e.g., high salinity, tide) can be, I also revealed significant outbreeding depression in reproductive traits. This helps explain how selfing is selectively maintained in rivulus, and along with other studies, helps us understand the temporal and spatially variable economics of mixed mating. That is, the interactive effects of treatment and heterozygosity on fecundity illustrates how the consequences of mixed mating change with environmental conditions. This emerging model system, rivulus, allows us to test hypotheses and provide critical insight about the maintenance of sex and sexual strategies previously only studied in plants and invertebrates

## SUPPORTING INFORMATION APPENDIX S1

**Table S1.1.** Experimental Lineage Summary. Includes the actual number of heterozygous microsatellites of the F0 progenitors, and the theoretical average number of heterozygous microsatellites for the experimental generation, the number of fish that survived to enter all treatments, the number that entered and survived each treatment, and the number that changed sex to male.

| Lineage | # Heterozygous microsatellites at F0 generation | # Heterozygous microsatellites at treatment generation | # Lived to enter treatment at 90dph | # Entered, # Finished CONTROL | # Entered, # Finished HIGH SALINITY | # Entered, # Finished LOW SALINITY | # Entered, # Finished TIDE | Number Changed Sex to Male |
| --- | --- | --- | --- | --- | --- | --- | --- | --- |
| BERM2 | 20 | 5 | 4 | 1,1 | 1,1 | 0,0 | 2,2 | 4 |
| BERM5 | 1 | 0 | 5 | 2,1 | 1,0 | 1,1 | 1,1 | 2 |
| BERM10 | 4 | 1 | 23 | 4,3 | 7,3 | 8,8 | 4,4 | 5 |
| BERM11 | 13 | 3 | 7 | 2,2 | 1,0 | 2,2 | 2,2 | 4 |
| BP1 | 6 | 2 | 4 | 1,1 | 0,0 | 3,2 | 0,0 | 1 |
| BP8 | 0 | 0 | 2 | 0,0 | 1,1 | 0,0 | 1,1 | 0 |
| BP9 | 4 | 0 | 20 | 5,5 | 4,4 | 7,6 | 4,4 | 1 |
| BP13 | 2 | 1 | 15 | 3,2 | 5,2 | 3,3 | 4,4 | 2 |
| BP15 | 1 | 0 | 44 | 11,9 | 12,5 | 10,6 | 11,9 | 5 |
| BP21 | 0 | 0 | 43 | 11,8 | 11,7 | 9,8 | 12,10 | 3 |
| BP23 | 0 | 0 | 24 | 6,6 | 6,4 | 3,3 | 9,5 | 14 |
| BUN1 | 0 | 0 | 32 | 4,4 | 8,4 | 9,9 | 11,6 | 7 |
| BUN2 | 0 | 0 | 2 | 0,0 | 1,1 | 1,1 | 0,0 | 0 |
| BUN3 | 0 | 0 | 11 | 2,2 | 2,1 | 3,3 | 4,2 | 4 |
| BUN5 | 0 | 0 | 22 | 5,5 | 5,3 | 6,5 | 6,6 | 1 |
| BUN6 | 0 | 0 | 6 | 3,2 | 0,0 | 3,3 | 0,0 | 1 |
| BUN7 | 0 | 0 | 9 | 4,4 | 2,1 | 1,1 | 2,2 | 1 |
| BWN1 | 17 | 4 | 8 | 0,0 | 2,1 | 1,1 | 5,4 | 5 |
| BWN2 | 21 | 6 | 8 | 4,4 | 1,0 | 1,1 | 2,2 | 4 |
| BWN3 | 0 | 0 | 9 | 3,3 | 1,1 | 3,3 | 2,1 | 4 |
| BWS11 | 0 | 0 | 20 | 3,3 | 7,3 | 5,5 | 5,3 | 2 |
| BWS21 | 2 | 1 | 14 | 4,4 | 1,0 | 4,4 | 5,5 | 0 |
| BWS30 | 7 | 2 | 15 | 2,2 | 4,1 | 3,2 | 6,5 | 3 |

| Lineage | # Heterozygous<br>microsatellites at<br>F0 generation | # Heterozygous<br>microsatellites at<br>treatment<br>generation | # Lived to enter<br>treatment at 90dph | # Entered, #<br>Finished<br>CONTROL | # Entered, #<br>Finished HIGH<br>SALINITY | # Entered, #<br>Finished LOW<br>SALINITY | # Entered, #<br>Finished TIDE | Number Changed<br>Sex to Male |
| --- | --- | --- | --- | --- | --- | --- | --- | --- |
| BWS32 | 3 | 1 | 13 | 4,3 | 3,2 | 4,3 | 2,2 | 5 |
| BWS37 | 3 | 1 | 29 | 6,5 | 10,4 | 7,7 | 6,6 | 0 |
| BWS38 | 7 | 2 | 14 | 3,3 | 6,2 | 3,3 | 2,2 | 1 |
| CF1-12 | 0 | 0 | 3 | 2,2 | 1,0 | 0,0 | 0,0 | 1 |
| CF4-18 | 0 | 0 | 1 | 0,0 | 0,0 | 0,0 | 1,1 | 0 |
| CF5-5 | 0 | 0 | 6 | 2,2 | 1,0 | 1,1 | 2,2 | 3 |
| CF15 | 0 | 0 | 9 | 3,3 | 2,2 | 3,3 | 1,1 | 4 |
| CROC21 | 18 | 5 | 3 | 0,0 | 0,0 | 2,2 | 1,1 | 0 |
| CROC22 | 2 | 1 | 14 | 4,4 | 2,0 | 4,3 | 4,1 | 1 |
| CROC23 | 6 | 2 | 11 | 1,1 | 3,3 | 4,4 | 3,3 | 4 |
| CROC29 | 18 | 5 | 1 | 0,0 | 1,1 | 0,0 | 0,0 | 0 |
| CROC31 | 20 | 5 | 4 | 2,2 | 1,1 | 0,0 | 1,0 | 0 |
| CRWL1 | 0 | 0 | 14 | 4,3 | 4,4 | 3,3 | 3,3 | 1 |
| CRWL15 | 0 | 0 | 23 | 4,3 | 5,2 | 6,6 | 8,8 | 1 |
| CRWL17 | 0 | 0 | 11 | 3,3 | 3,2 | 4,4 | 1,1 | 3 |
| CRWL19 | 0 | 0 | 23 | 5,5 | 8,4 | 5,4 | 5,5 | 2 |
| DC6 | 2 | 1 | 5 | 0,0 | 3,1 | 2,2 | 0,0 | 1 |
| DC8 | 16 | 4 | 8 | 1,1 | 2,1 | 3,3 | 2,2 | 2 |
| DC17 | 3 | 1 | 7 | 1,1 | 2,2 | 1,1 | 3,3 | 1 |
| DC21 | 0 | 0 | 5 | 0,0 | 2,0 | 2,2 | 1,1 | 1 |
| LION2 | 0 | 0 | 4 | 0,0 | 1,1 | 1,1 | 2,2 | 1 |
| LION5 | 0 | 0 | 1 | 0,0 | 1,1 | 0,0 | 0,0 | 0 |
| LK13 | 0 | 0 | 2 | 0,0 | 1,0 | 0,0 | 1,1 | 0 |
| LK15 | 0 | 0 | 10 | 1,1 | 4,2 | 4,3 | 1,1 | 4 |
| LMC1 | 3 | 1 | 13 | 3,2 | 3,1 | 4,3 | 3,3 | 6 |
| LMC3 | 0 | 0 | 11 | 3,3 | 1,0 | 3,3 | 4,3 | 0 |
| LMC4 | 4 | 1 | 21 | 4,4 | 6,1 | 3,3 | 8,6 | 10 |
| NC18 | 6 | 2 | 6 | 1,1 | 2,0 | 2,1 | 1,1 | 2 |
| NC19 | 4 | 1 | 10 | 1,1 | 4,2 | 3,2 | 2,2 | 4 |
| NC23 | 10 | 3 | 4 | 0,0 | 0,0 | 1,0 | 3,3 | 1 |
| NNKB1 | 0 | 0 | 21 | 6,6 | 5,4 | 5,5 | 5,4 | 4 |
| NNKB3 | 8 | 1 | 1 | 1,1 | 0,0 | 0,0 | 0,0 | 0 |
| PLT2 | 4 | 1 | 1 | 0,0 | 0,0 | 1,1 | 0,0 | 0 |
| SAND19 | 7 | 2 | 15 | 4,3 | 4,1 | 3,2 | 4,3 | 0 |
| SAND21 | 2 | 1 | 15 | 3,3 | 6,3 | 3,2 | 3,3 | 2 |
| SAND24 | 17 | 5 | 24 | 6,5 | 8,3 | 5,4 | 5,5 | 7 |
| SAND29 | 0 | 0 | 19 | 4,4 | 5,4 | 5,3 | 5,5 | 6 |
| SAND36 | 4 | 1 | 5 | 1,1 | 2,0 | 1,1 | 1,1 | 0 |
| SAND38 | 8 | 2 | 9 | 2,2 | 2,0 | 4,2 | 1,1 | 4 |
| SAND40 | 15 | 4 | 13 | 3,3 | 6,1 | 2,2 | 2,2 | 1 |
| SOB6 | 0 | 0 | 13 | 4,4 | 3,3 | 3,3 | 3,2 | 6 |
| SOB7 | 5 | 1 | 17 | 5,3 | 4,1 | 3,2 | 5,4 | 6 |
| SOB9 | 3 | 1 | 33 | 6,6 | 7,2 | 11,9 | 9,8 | 4 |
| UM2 | 2 | 1 | 18 | 3,3 | 4,2 | 7,6 | 4,4 | 4 |
| UM3 | 0 | 0 | 20 | 5,4 | 4,4 | 4,4 | 7,4 | 1 |
| UM9 | 1 | 0 | 13 | 4,4 | 2,2 | 4,4 | 3,3 | 3 |

**Table S1.2.** Summary of regression analysis models for size traits. All juveniles were kept in the same constant condition of 25‰. There were four treatments in the treatment period (control, low salinity, high salinity, and tide). Heterozygosity was modeled as a continuous variable ranging from 1 - 6 heterozygous microsatellites (out of 32). Significant effects are shown in bold. F = F-ratio; P = P-value; df = degrees of freedom.

| Model Variables |  | Mass |  |  | Length |  |  |  | Fulton's K Body Condition |  |  |  |
| --- | --- | --- | --- | --- | --- | --- | --- | --- | --- | --- | --- | --- |
| Juvenile Period (90dph) | Estimate | F | df | P | Estimate | F | df | P | Estimate | F | df | P |
| Heterozygosity | 0.0008 | 3.35 | 1, 866 | 0.07 | **0.099** | **7.28** | **1, 861** | **0.01** | -0.006 | 2.15 | 1, 857 | 0.14 |
|  |  | Mass |  |  | Length |  |  |  | Fulton's K Body Condition |  |  |  |
| Treatment Period (180dph) | Estimate | F | df | P | Estimate | F | df | P | Estimate | F | df | P |
| Treatment |  | **294.66** | **3, 3.50** | **< 0.0001** |  | **85.59** | **3, 4.84** | **< 0.0001** |  | **41.57** | **3, 3.69** | **0.003** |
| Heterozygosity | -0.0007 | 0.54 | 1, 647.6 | 0.46 | 0.011 | 0.05 | 1, 658.2 | 0.83 | **-0.011** | **4.64** | **1, 647.3** | **0.03** |
| Treatment x Heterozygosity |  | 0.05 | 3, 630.4 | 0.99 |  | 0.09 | 3, 657.2 | 0.96 |  | 1.29 | 3, 636.8 | 0.28 |

**Figure S1.1.**
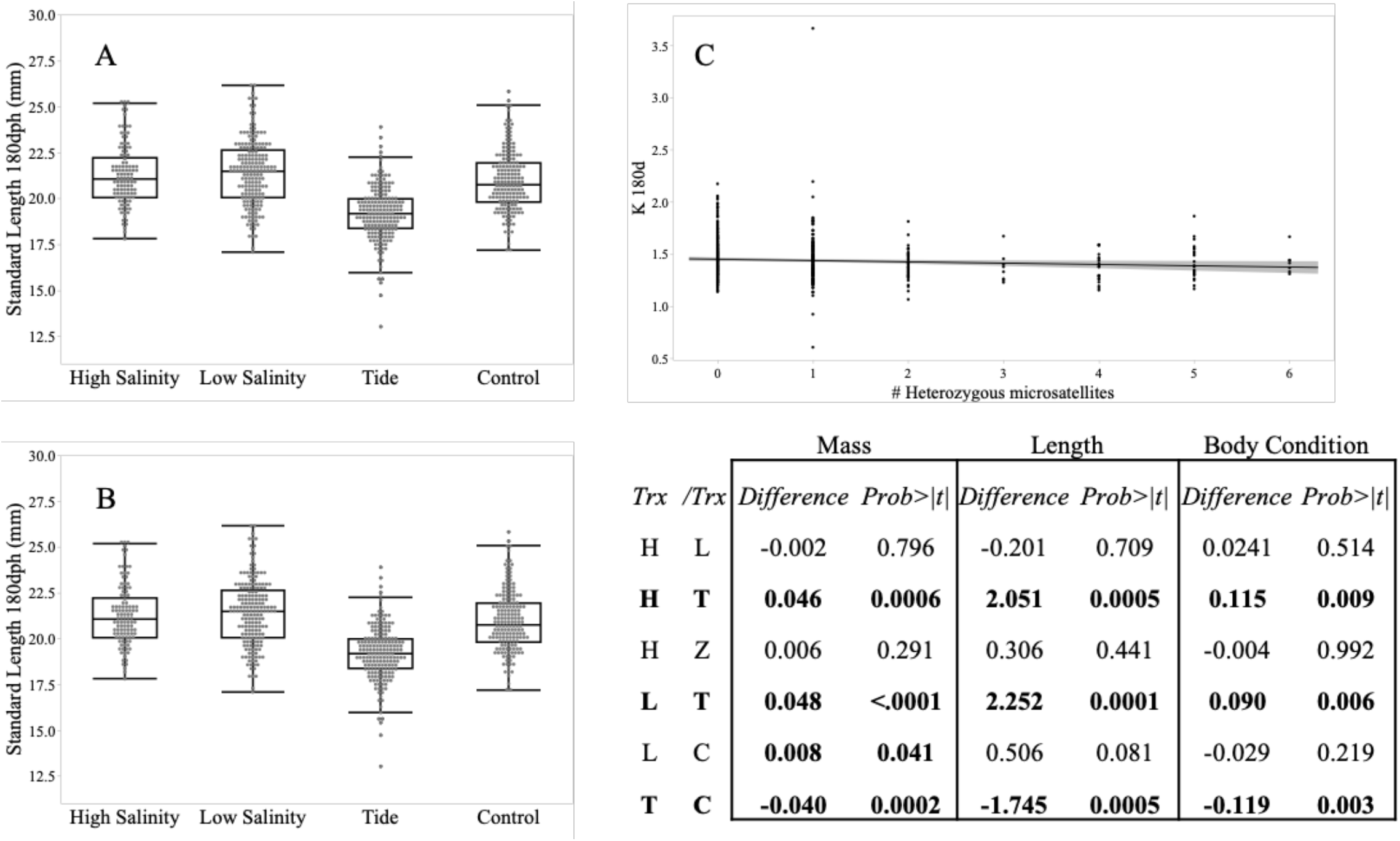
Treatment differences in two size traits (A. standard length and B. body condition, mass is Figure 2 within the text), when measured at the end of the treatment period (180 dph). Those fish in the tide treatment were significantly smaller in mass, standard length, and body condition than those in all conditions. Those fish in the low salinity also had a significantly larger mass than those fish in the control treatment. C. Body condition decreases with increasing heterozygosity in all treatments. Significant effects are shown in bold. P = P-value.

**Figure S1.2.**
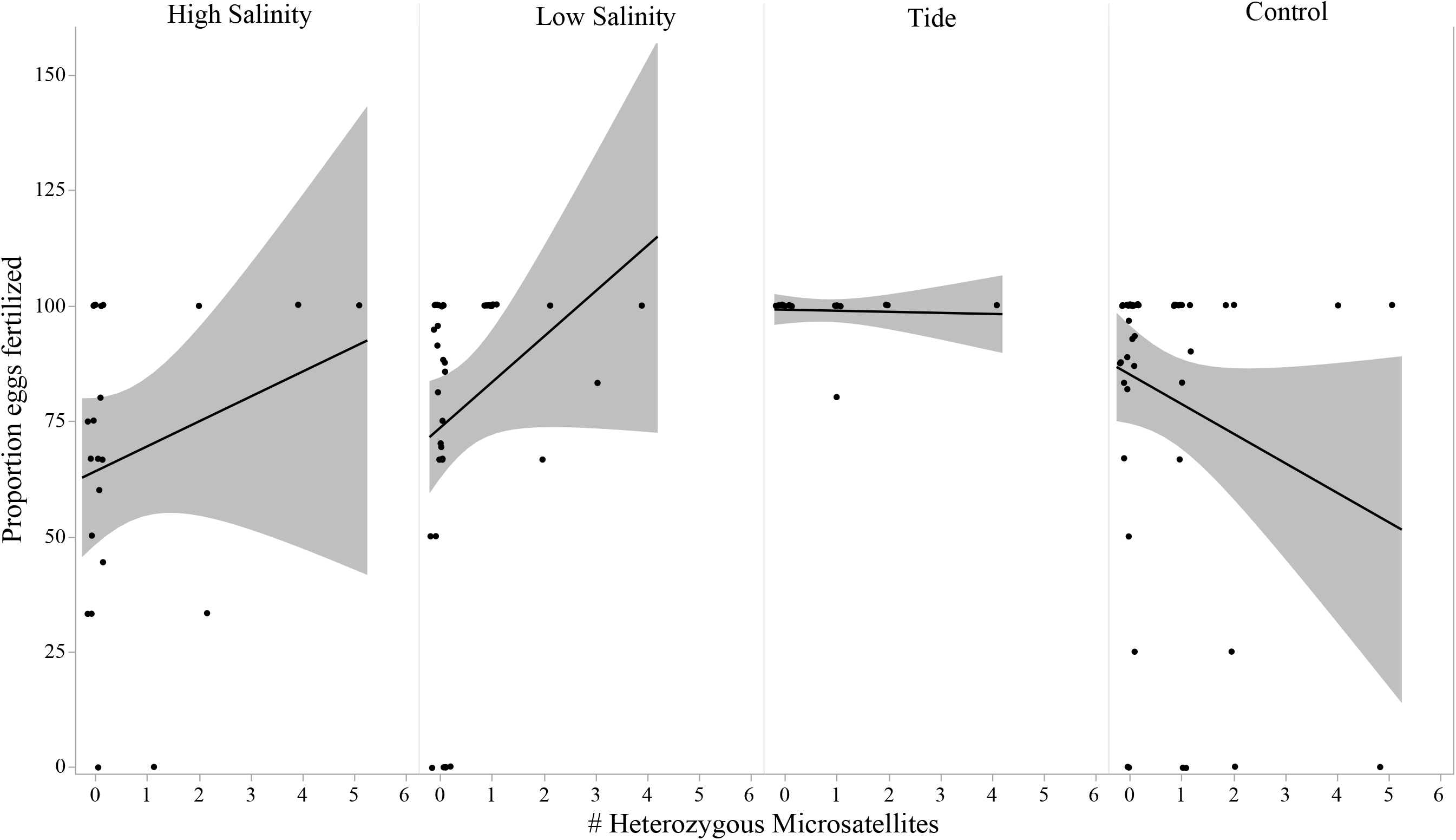
Treatment and heterozygosity effects on the proportion of fertilized eggs. Proportion significantly decreased with increasing heterozygosity in the control treatment, while the inverse is significant in the high salinity and low salinity treatments. There was not an effect of heterozygosity in the tidal treatment as almost all eggs were fertilized.

**Table S1.3:** Treatment (Trx) effects on whether or not individual fish laid any eggs. Treatment contrasts with significant differences shown in bold. Those fish in the tide and high salinity treatments were less likely to lay eggs than those in control and low salinity. Significant comparisons are shown in bold. SE = standard error; P = P-value.

| *Treatment contrasts for Binomial Response of Laying Eggs* |  |  |  |  |  |
| --- | --- | --- | --- | --- | --- |
| *Trx* | */Trx* | *Odds Ratio* | *SE* | *Z-ratio* | *P* |
| **C** | **H** | **4.20** | **1.30** | **4.64** | **<.0001** |
| C | L | 1.63 | 0.46 | 1.74 | 0.31 |
| **C** | **T** | **4.67** | **1.51** | **4.76** | **<.0001** |
| **H** | **L** | **0.39** | **0.13** | **-2.95** | **0.02** |
| H | T | 1.11 | 0.40 | 0.29 | 0.99 |
| **L** | **T** | **2.86** | **0.96** | **3.14** | **0.01** |

**Table S1.4.**
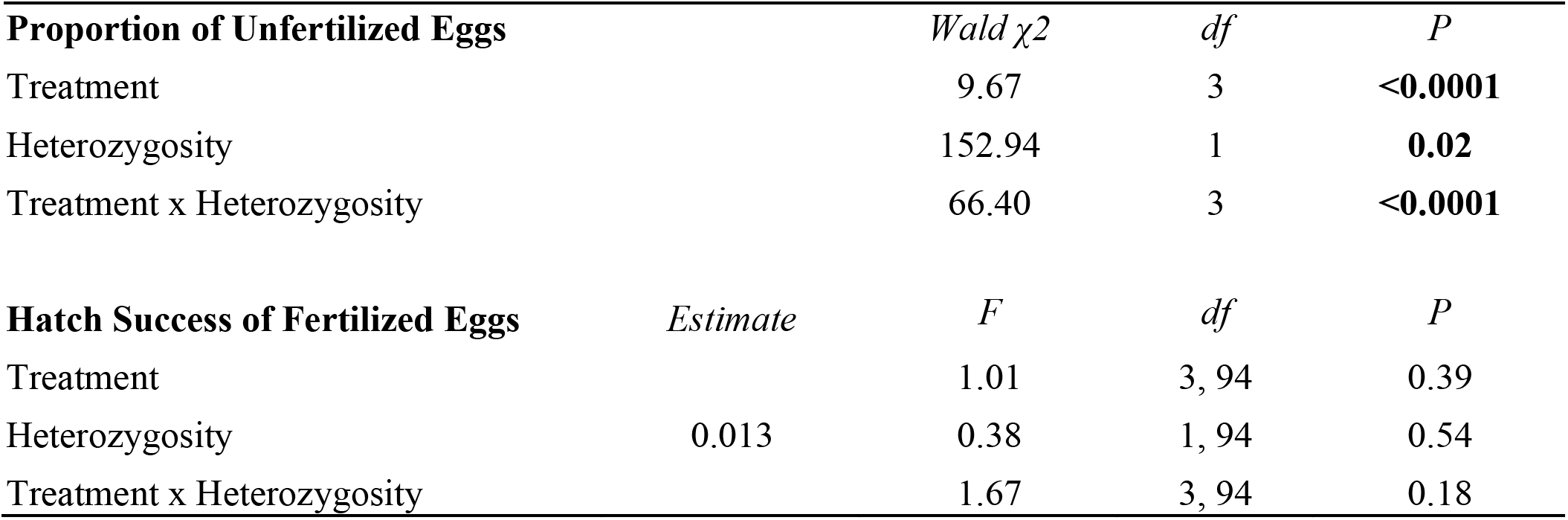
Treatment and heterozygosity effects on the proportion of unfertilized eggs and hatch success of fertilized eggs. Significant values are shown in bold. There was an interaction of treatment and heterozygosity on the proportion of unfertilized (and thus fertilized) eggs. See Figure S2. Hatch success of fertilized eggs was not affected by treatment or heterozygosity. F = F-ratio, P = p-value, df=degrees of freedom.

**Table S1.5.**
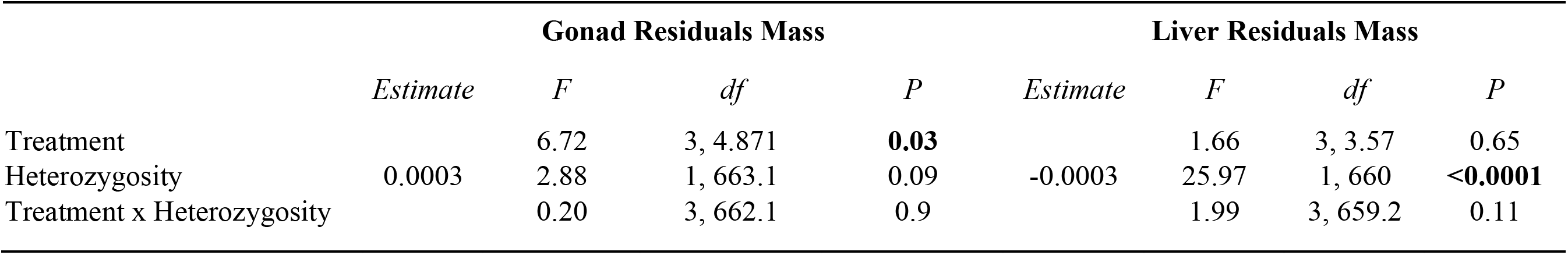
Treatment and heterozygosity effects on the gonad and liver masses, relative to body mass. Significant values are shown in bold. Only treatment affected gonad mass. Gonad mass was significantly larger in fish in the control treatment compared to those in the low salinity treatment. There were not differences in other pairwise comparisons. Liver mass was only affected by heterozygosity. Liver mass decreased as heterozygosity increased. F = F-ratio, P = p-value, df=degrees of freedom.

**Table S1.6.** Treatment and size effects on survival until the end of treatment at 180 dph. Each size trait, plus treatment and the interaction of the size trait and treatment, were run as separate models. The range of Wald χ^2^ values for treatment reflect the values from the different models. Survival decreased with decreasing mass and standard length at 90 dph and 150 dph, the negative parameter estimate indicates that the hazard of mortality decreases with the increase of the independent variable. Body condition was not a significant effect. Treatment alone was a significant effect. F = F-ratio, P = p-value, df=degrees of freedom.

| **Effects of Size on Survival to 180 dph** | *Estimate* | *Wald $\chi^2$* | *df* | *P* |
| --- | --- | --- | --- | --- |
| 90 dph mass (mg) | **-0.012** | **8.25** | **1** | **0.004** |
| 90 dph standard length (mm) | **-0.152** | **9.09** | **1** | **0.003** |
| 90 dph body condition (Fulton's K) | 0.114 | 0.07 | 1 | 0.790 |
| **Treatment** |  | **22.15-23.52** | **1** | **<0.0001** |
| 150 dph mass (mg) | **-0.030** | **8.84** | **1** | **0.003** |
| 150 dph standard length (mm) | **-0.352** | **6.75** | **1** | **0.009** |
| 150 dph body condition (Fulton's K) | -1.747 | 1.76 | 1 | 0.185 |
| 150 dph mass (mg) x treatment |  | 2.30 | 3 | 0.513 |
| 150 dph standard length (mm) x treatment |  | 2.19 | 3 | 0.534 |
| 150 dph body condition (Fulton's K) x treatment |  | 1.95 | 3 | 0.583 |

